# MHC heterozygosity may increase subordinate but not alpha male siring success in white-faced capuchin monkeys (*Cebus imitator*)

**DOI:** 10.1101/2024.09.04.611202

**Authors:** Janet C. Buckner, Katharine M. Jack, Margaret Buehler, Amanda D. Melin, Valérie A. M. Schoof, Eva C. Wikberg, Saul Cheves Hernandez, Linda M. Fedigan, Jessica W. Lynch

## Abstract

The genes of the major histocompatibility complex (MHC) are vital to vertebrate immunity and may influence mate choice in several species. The extent to which the MHC influences female mate choice in primates remains poorly understood, and studies of MHC-based mate choice in platyrrhines are especially rare. White-faced capuchin monkeys (*Cebus imitator*) reside in multimale-multifemale groups where alpha males sire most of the offspring. In this study, we investigated the roles of social dominance, relatedness, and MHC genotypes in determining which mating pairs produced offspring in wild white-faced capuchins in the Sector Santa Rosa (SSR), Área de Conservación Guanacaste, Costa Rica. We find that males in this population do not differ significantly in MHC metrics based on their social status or siring success. Using mixed conditional logit models and generalized linear models, we find that alpha males that are distantly related to reproducing females are significantly more likely to sire offspring while MHC metrics do not predict the probability of siring offspring, or becoming an alpha male. However, we do find some evidence that subordinate males heterozygous at MHC loci sire significantly more offspring than homozygous subordinates. Further, one-sided binomial simulations reveal that offspring are more frequently heterozygous at MHC loci than expected given the gene pool. We conclude that in this population with limited genomic variation, females may preferentially mate with MHC-diverse subordinate males when related to the alpha, leading to increased probabilities of MHC-diverse offspring.

## Introduction

A longstanding hypothesis in sexual selection theory posits that individuals, typically females, key into cues that signal “good genes” in a potential mate that can be passed to offspring. Alternatively, females choose mates based on genetic dissimilarity, or “compatible genes”, as maximizing genetic diversity in offspring may increase fitness. These distinct hypotheses, which potentially operate simultaneously, have garnered notable interest from the scientific community (Kirkpatrick, 1987; Tregenza and Wedell, 2001; Mays Jr. and Hill, 2004; Neff and Pitcher, 2005; Puurtinen et al., 2009; Achorn and Rosenthal, 2020). The genes of the major histocompatibility complex (MHC) have often been at the center of this controversial area of inquiry into the genetic basis of mate choice.

The MHC is a group of highly polymorphic genes primarily responsible for encoding proteins vital for the function of adaptive and innate immune responses (Janeway Jr et al., 2001). Increased variation in MHC alleles allows individuals to recognize and combat a greater range of potential intracellular and extracellular pathogens. Additionally, these genes may function in kin recognition and inbreeding avoidance as the MHC helps the body discriminate between self and non-self (Brown and Eklund, 1994; Adams and Parham, 2001). Thus, MHC genotypes can impact individual survivorship, mating decisions, and ultimately fitness. If individuals can discern MHC genotypes, these may provide honest information about the potential of different mates to maximize offspring survival and fitness. While support for MHC-mediated mate choice is inconsistent in non-human primates, several studies have presented evidence of non-random mating associated with MHC genotypes, related to the *disassortative mate choice* and *diversity-advantaged mate choice* hypotheses (reviewed in Setchell and Huchard, 2010; Huchard and Pechouskova, 2014, Kamiya et. al., 2014).

The *disassortative mate choice* hypothesis proposes that mates are selected to maximize their MHC dissimilarity with the chooser, which provides fitness benefits such as inbreeding avoidance and increased pathogen resistance in offspring due to increased MHC diversity. Evidence for disassortative mating has been found in various non-human primates (grey mouse lemurs [*Microcebus murinus*] – Schwensow et al., 2008b, Huchard et al., 2013; mandrills [*Mandrillus sphinx*] - Setchell et al., 2016; golden snub-nosed monkeys [*Rhinopithecus roxellana*] - Zhang et al., 2020).

The *diversity-advantaged mate choice hypothesis* (MHC-diverse mate choice herein) proposes that mates are selected because they possess diverse MHC alleles (e.g., heterozygotes). Theoretically, heterozygotes possess more rare alleles than homozygotes on average, thus increasing the likelihood that they will pass on advantageous rare alleles. A meta-analysis recovered a trend of preference for MHC-diverse mates in non-human primates across six species (Winternitz et al., 2017), although some studies of the same species did not find this preference. In the only study to date on the influence of MHC mate choice in a platyrrhine primate, the northern muriqui [*Brachyteles hypoxanthus]* showed diversity-advantaged mate choice but not disassortative mate choice (Chaves et al., 2023). Strategies for MHC-mediated mate choice are not mutually exclusive and may operate simultaneously, or under different conditions, in the same species or population.

When investigating mate choice, it is important to consider social structure, and particularly the effect that dominance relationships can have on limiting the expression of female mate choice. Even within multi-female multi-male groups, primate social structures can range from hierarchical societies dominated by single males that sire most of the offspring (high reproductive skew), to egalitarian or female-dominated societies with more opportunity for females to express mate choice. In other words, the relationship between MHC variability and female mate choice may be stronger or weaker depending on the socio-ecologically determined mating system. In muriquis, a multimale-multifemale egalitarian society, there was evidence for MHC-diverse mate preferences (Chaves et al., 2023). Evidence for both disassortative mating and MHC-diverse female mate preferences were found in mandrills living in a multimale-multifemale group with high reproductive skew towards the alpha male. However this study was performed on a closed semi-free ranging population with 15 founders, and thus included human influence on the group composition (Setchell et al., 2009). Some other studies of multimale-multifemale groups, including golden snub-nosed monkeys, chacma baboons [*Papio ursinus*], and rhesus macaques [*Macaca mulatta*] also found evidence for MHC-diverse mate preferences, even though dominance hierarchies are fairly rigid and males are dominant to females in these species (Winternitz et al., 2017).

Accordingly, there may be mechanisms by which females can exert mate choice even in male-dominant societies with rigid hierarchies and high reproductive skew. For example, females could influence which males enter their social group and which males rise to alpha status. They could also mate selectively with different males depending on their fertility status (Carnegie et al., 2006; Manson et al., 1997). Females of some species have been shown to exert post-copulatory gamete choice, increasing the likelihood of producing a heterozygous offspring (Engelhardt et al., 2006; Schwensow et al., 2008a). MHC molecules are expressed on the surface of spermatozoid (Paradisi et al., 2000) and studies of mice have shown that their oocytes can differentially select sperm with dissimilar MHC haplotypes (Wedekind et al., 1996; Rűlicke et al., 1998). Alcaide et al. (2012) suggests that a similar selection for “*genetically loaded spermatozoa*” may be occurring in female kestrels [*Falco naumanni*]. However, in the only known study testing this in non-human primates, Setchell and colleagues (2013) found no evidence for post-copulatory selection for MHC genotypes in mandrills.

It is also likely that if the MHC does play a role in mate choice for a particular species, its function (i.e., the basis of the mate choice) may fluctuate with changing environmental pressures (e.g., changes in parasite and pathogen populations or possibly even population size) (e.g., Evans et al., 2012). This may explain studies of non-human primates where support for MHC-mediated mate choice is inconsistent (golden snub-nosed monkeys - Yang et al., 2014; Zhang et al., 2020) or completely absent (e.g., chacma baboons - Huchard et. al., 2010). Collectively, these studies indicate that MHC-based mate choice likely functions in different ways for different species or populations (Piertney and Oliver, 2006).

In this study, we investigated the roles of social dominance, relatedness, and MHC genotypes in determining which pairs produced offspring in wild white-faced capuchins [*Cebus imitator*] in the Sector Santa Rosa (SSR), Área de Conservación Guanacaste, Costa Rica. The SSR capuchins reside in multimale-multifemale groups (mean = 15.9 group members), and alpha males sire most of the offspring (high reproductive skew) (Wikberg et al., 2017; Muniz et al., 2010; Jack and Fedigan, 2006). Populations also exhibit low local genetic diversity due to the social organization and dispersal strategies (Orkin et al., 2021). Females are philopatric and closely related, with new groups generally formed when large groups fission. Males first disperse at approximately four years of age and can change groups throughout their lives (Jack and Fedigan, 2004a, 2004b; Jack et al., 2012). However, males usually disperse in cohorts or selectively join neighboring groups (Jack and Fedigan, 2004b), which decreases the genetic diversity of males in an area, so that immigrant males residing within groups at our SSR are often as closely related as natal females (Wikberg et al., 2014). Studies have shown that MHC can be considerably diverse despite such low overall diversity in the rest of the genome (Oliver & Piertney, 2012; Mona et al., 2008; Van Oosterhout et. al., 2006; Aguilar et al., 2004), and MHC-mediated mate choice may be a mechanism that leads to such observed patterns. In fact, some nonhuman primate studies indicate that selective pressures for MHC disassortative mating are highest in populations with limited outbreeding opportunities (Roberts et al., 2010a). While female mate choice opportunities seem low in this male-dominated hierarchical social system, female white-faced capuchins may be able to exert mate choice at the behavioral level by influencing male dominance rank within groups (Perry, 1998) and selectively mating with preferred males during fertile periods (Carnegie et al., 2006), as well as potentially at the physiological level, through post-copulatory sperm selection (see above).

The capuchins at Santa Rosa have a long life span (>25 years for both males and females), interact with large numbers of individuals, have an omnivorous high energy diet with destructive foraging (Melin et al., 2014; Bergstrom et al., 2018), high parasite loads (Parr et. al., 2013) and are non-seasonal breeders (Carnegie et al., 2011) that engage in non-conceptive copulations (Manson et al., 1997). We leveraged behavioral and genetic data to analyze multiple metrics of MHC variability in parental-offspring triads over 18 years of study (monitoring births from 1996 to 2014) to test for signals of MHC-mediated female mate choice. Given the high degree of sociality, and the ecological and genetic challenges to this population, we predicted that there would be strong selection for a) high MHC diversity in alpha males (*diversity-advantaged mate choice hypothesis)* and/or b) females will show preferences for pairings between MHC disparate partners to maximize MHC allele diversity in offspring (disassortative mate choice hypothesis).

## Methods

### Ethics statement

Our research followed the American Society of Primatologists (ASP) Principles for the Ethical Treatment of Non-Human Primates. We collected samples in accordance with the laws of Costa Rica, the United States, and Canada and complied with protocols approved by the Área de Conservación Guanacaste and by Tulane University’s Animal Care and Use Committee (Protocol 0399R2) and the Canada Research Council for Animal Care through the University of Calgary’s Life and Environmental Care Committee (ACC protocols AC15-0161 and AC20-0148). We obtained permits for sample collection from the Área de Conservación Guanacaste (ACG-PI-033-2016 and PI-063-2014) and CONAGEBIO (R-001-2015-OT-CONAGEBIO and R-025-2014-OT-CONAGEBIO), sample export from Costa Rica under permits from the Área de Conservación Guanacaste (DGVS-030-2016-ACG-PI-002-2016; 012706), and imported to Canada with permission from the Canadian Food and Inspection Agency (A-2016-03992-4) and to the United States with permission from the Center for Disease Control (PHS Permit No. 2010-01-140).

### Population and Demographics

This study focuses on a single wild population of white-faced capuchin monkeys (*Cebus imitator*) from Sector Santa Rosa (SSR) of the Área de Conservación Guanacaste, Costa Rica. Within SSR, there are approximately 48 groups with an average size of 15.9 individuals (Hogan et al., 2019). The primates in this population have been under near continuous study for four decades resulting in extensive data on social structure, dominance, mating behavior, parentage, dispersal, and population genetics. We specifically leverage neutral genetic data and pedigree information to evaluate MHC diversity and reproductive outcomes in a sample of 115 individuals across four groups. Fecal samples used for this study were collected between 2005 and 2013, with the earliest sample coming from a female born in 1994. This data set required the analysis of samples from infants, mothers, and potential sires, which could not be done for all infants born during the study period due to the deaths or disappearances of potential sires and/or mothers. In some cases we were able to infer MHC genotypes from an infant with a known maternal genotype based on information gained through the analysis of siblings. Our data set thus includes 95 infants born to 34 different females. Of the 45 adult and subadult males included in the current study, 20 sired offspring (44%). Alpha males sired 69 offspring (73%), while subordinates sired the remaining 26 infants (27%). These infants were sired by 8 different alpha males, 9 different subordinate males, and 3 additional males that sired offspring as both an alpha and as a subordinate.

### Genotyping

Multiple fecal samples (2-5 samples/individual) were collected from each study subject. We placed 1 gm of fresh feces collected from an identified monkey by an experienced researcher immediately in 5ml of RNAlater. DNA was extracted from fecal samples using a Macherey-Nagel NucleoSpin Tissue kit, following manufacturer’s instructions. Parentage for individuals included in the study was determined as part of Wikberg et al. (2014; 2017). Briefly, paternity and estimated relatedness values were determined via genotyping from fecal DNA of up to 20 Short Tandem Repeat loci. The MHC genotype data used in this study were sequenced and processed as part of Buckner et al. (2021). Briefly, four MHC Class II DR and DQ exons were genotyped from 115 individuals using non-invasive sampling and high throughput sequencing. We confirmed haplotypes and genotypes by using the pedigree information available for the study population. In some cases, poor sample quality (i.e., degraded DNA from fecal samples) precluded genotyping of one or more exons for some individuals. As MHC alleles are typically inherited as a linked set from each parent (Knapp, 2005), we were able to use parent-offspring triad information to confirm haplotypes and fill in missing exon data when individuals had partial data available. We were not able to sequence MHC exons for all of the potential sires and infants during the span of the study years. Our MHC data set included 39 parent-offspring triads, and we included MHC data for 28 potentially reproducing males in the population.

Here, we focused our analyses on MHC-DRB exon 2 genotypes as they are considered to display the highest degree of polymorphism within MHC across humans and several non-human primates (de Groot et. al., 2012). The antigen binding site, which binds and presents foreign antigens to T-cells, is encoded by exon two in Class II MHC genes and as a result MHC-DRB exon 2 is the most widely studied exon across mammalian species. Therefore, it may justifiably serve as a reduced representation of the overall MHC diversity, and we report our results from analyzing the MHC-DRB exon 2 below. However, different MHC genes can show different signatures of mate choice (Huchard et. al., 2013); therefore, we performed analyses on all four sequenced exons (DRB exon 2, DRB exon 3, DQA exon 3, DQB exon 3) separately and perform a set of analyses on all four exons jointly (Supplemental Information).

### Statistical analyses

Following the work by Santos et al. (2016) and Setchell et al. (2010) on MHC-mediated mate choice, we determined the evidence for MHC-disassortative and MHC-diverse mate choice by calculating a set of seven metrics of MHC variability across males and mate pairs (Table 1). We compared two aspects of MHC genotypes; the unique alleles present in each individual and the diversity of the amino acids produced by those alleles. For each individual, we determined which unique alleles were present at each of the 4 exons (exon 3 of DQA, DQB and DRB, and DRB exon 2) as well as the number of differences between the amino acids produced by each pair of alleles. Using these data, we calculated diversity and dissimilarity metrics (Table 1; see Santos et al., 2016; Setchell et al., 2010). Diversity metrics (suffix Div) refer to the number of unique alleles or amino acids *within* a single individual. Dissimilarity metrics (suffix Dis) compare alleles or amino acids *between* a pair of individuals. Finally, we tested for correlations between MHC metrics and included only sets of metrics with pairwise R values <0.80 in the models.

**Table 1.**
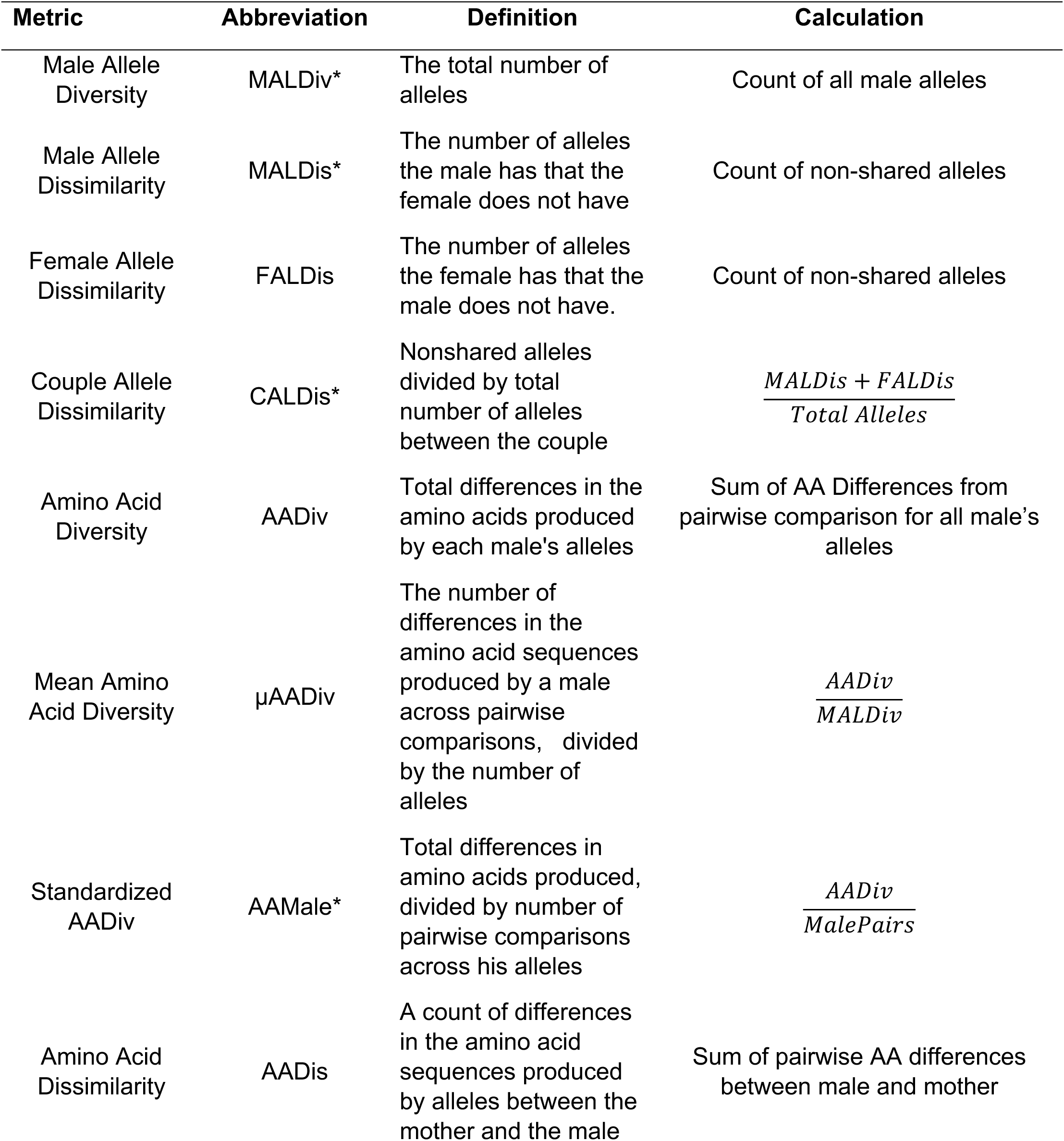
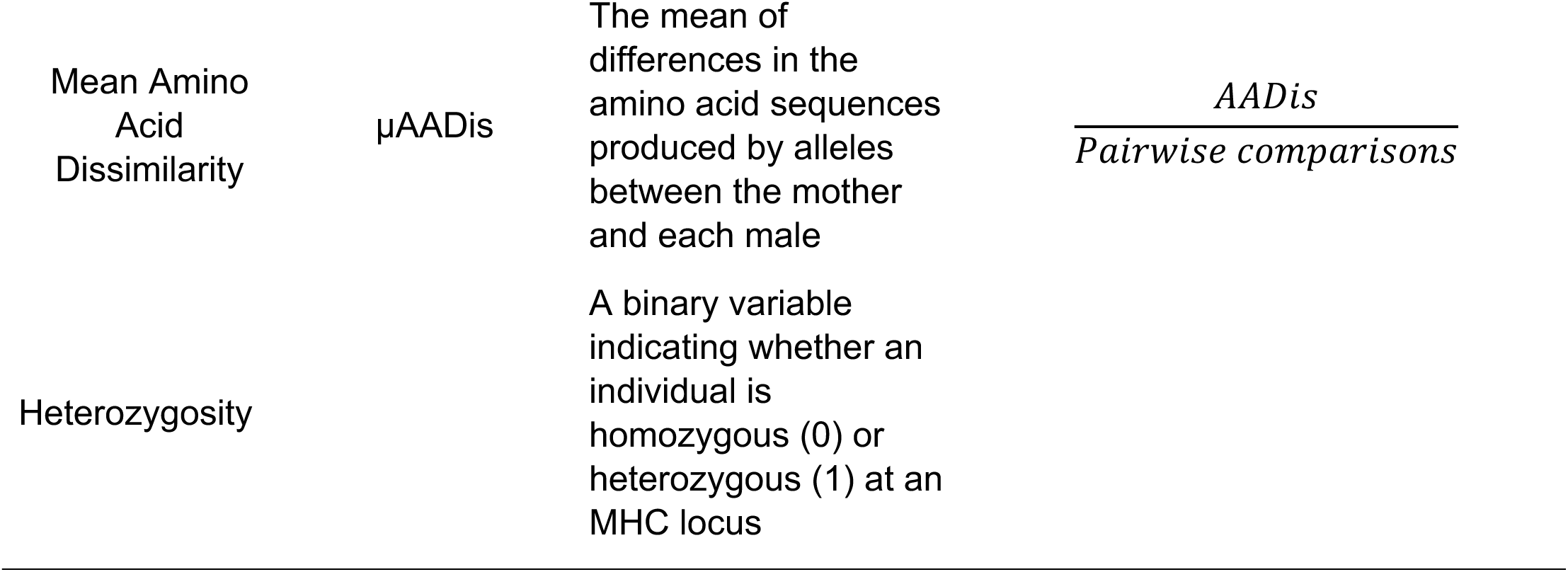
Abbreviations, definitions and calculations for MHC metrics used in this study. Asterisks (*) indicate variables that were included in the t-tests, glm and/or mclogit analyses.

| **Metric** | **Abbreviation** | **Definition** | **Calculation** |
| --- | --- | --- | --- |
| Male Allele Diversity | MALDiv* | The total number of alleles | Count of all male alleles |
| Male Allele Dissimilarity | MALDis* | The number of alleles the male has that the female does not have | Count of non-shared alleles |
| Female Allele Dissimilarity | FALDis | The number of alleles the female has that the male does not have. | Count of non-shared alleles |
| Couple Allele Dissimilarity | CALDis* | Nonshared alleles divided by total number of alleles between the couple | $\frac{MALDis + FALDis}{Total\ Alleles}$ |
| Amino Acid Diversity | AADiv | Total differences in the amino acids produced by each male's alleles | Sum of AA Differences from pairwise comparison for all male's alleles |
| Mean Amino Acid Diversity | μAADiv | The number of differences in the amino acid sequences produced by a male across pairwise comparisons, divided by the number of alleles | $\frac{AADiv}{MALDiv}$ |
| Standardized AADiv | AAMale* | Total differences in amino acids produced, divided by number of pairwise comparisons across his alleles | $\frac{AADiv}{MalePairs}$ |
| Amino Acid Dissimilarity | AADis | A count of differences in the amino acid sequences produced by alleles between the mother and the male | Sum of pairwise AA differences between male and mother |
| Mean Amino<br>Acid<br>Dissimilarity | μAADis | The mean of<br>differences in the<br>amino acid sequences<br>produced by alleles<br>between the mother<br>and each male | *AADis*<br>*Pairwise comparisons* |
| Heterozygosity |  | A binary variable<br>indicating whether an<br>individual is<br>homozygous (0) or<br>heterozygous (1) at an<br>MHC locus |  |

We considered each infant sired to be a single conception event in which a female chose a specific male to sire her offspring. We excluded cases in which there was only one male present in the group at conception (N=3) as this precluded female choice. For each conception event, we recorded each male’s social status in the group (alpha or subordinate), the relatedness values between the mother and each male, as well as each individual’s MHC genotype. We structured our dataset so that each male present in the group at the time of siring was listed as a sire or non-sire for each offspring, and thus, males are evaluated each time a female in their group conceived an infant. This repetition is representative, as each instance of female mate choice depends upon the options available in the group at the time. While a male may have been a potential mate at the conception of some of a female’s offspring, her options could differ depending on which other males were in the group at the time and which male held alpha status. Some males are present in the dataset as a subordinate male and as an alpha male at different time points. While a male’s genotype will not change between two opportunities to sire an offspring, his social rank, his group membership, and/or his male competitors in the group could, all of which might impact female mate choice. Each male’s dissimilarity statistics (MALDis, CALDis, AADis, and µAADis) also differ depending on the genotype of the female.

We ran analyses on two separate datasets: (1) each MHC metric calculated separately for each of the four exons and (2) the sum of each MHC metric across all four exons. The first dataset allowed us to evaluate if specific exons had a greater influence on female choice than others. We first ran a series of T-tests to determine if the mean value of each diversity metric differed significantly between sires and non-sires or between alpha and subordinate males. We fit mixed conditional logit models using the R package *mclogit* (Elff, 2009; Elff, 2022) with male ID as a random effect and relatedness, rank, and MHC metrics as fixed effects to test if the latter variables significantly predict the likelihood for a male to sire offspring. We also fit *mclogit* models that included only offspring sired by subordinate males for the MHC dataset as prior data suggested mate choice behavior may change when the alpha is a relative. Before running the analysis, we tested for pairwise correlations between each MHC metric using the base package in R, and removed one of the two metrics when they were significantly correlated with an R^2^ ≥ 0.80. We also fit generalized linear models (glm) using the base *stats* package in R with family specified as “binomial” to test if MHC metrics could predict the likelihood of a male attaining alpha status. Variables were considered significant if the p-value was less than or equal to 0.05.

We also tested whether pre-copulatory female mate choice and/or post-copulatory gamete selection favors MHC heterozygosity in offspring in the Santa Rosa capuchin population. First, we considered the overall probability of a heterozygous offspring given the males available to the female for each conception event, averaging across conception events to calculate a population-wide ‘expected MHC heterozygosity’ for offspring. Given the genotypes of parental pairs for each conception event, we then calculated the expected outcomes in terms of heterozygous versus homozygous MHC for resultant offspring using Mendelian genetics. Then we compared the expected population-wide and pair-specific outcomes to the observed genotypes in these offspring in our birth sample. Finally, we ran one-sided binomial simulations with 1,000 iterations based on the expected probability for heterozygotes given the mating pool, and the identified sire, to determine if the observed proportion of heterozygous offspring was higher than expected by chance.

## Results

### Alphas, subordinates, sires, and non-sires do not differ significantly in mean MHC metrics

The results from the statistical tests on DRB exon 2 are summarized in Tables 2-4 and Figure 1. Data from the remaining three exons and combined dataset are reported in Tables S1-S3. When evaluating mean values of MHC diversity metrics across males, there were no significant differences between sires and non-sires or alphas and subordinates (Table 2). However, AADiv, AAMale and uAADiv at DRB exon 3 as well as AAMale and MALDiv at DQA exon 3 displayed nearly significant differences (<0.1) between alpha males and subordinates (Table S1).

**Figure 1.**
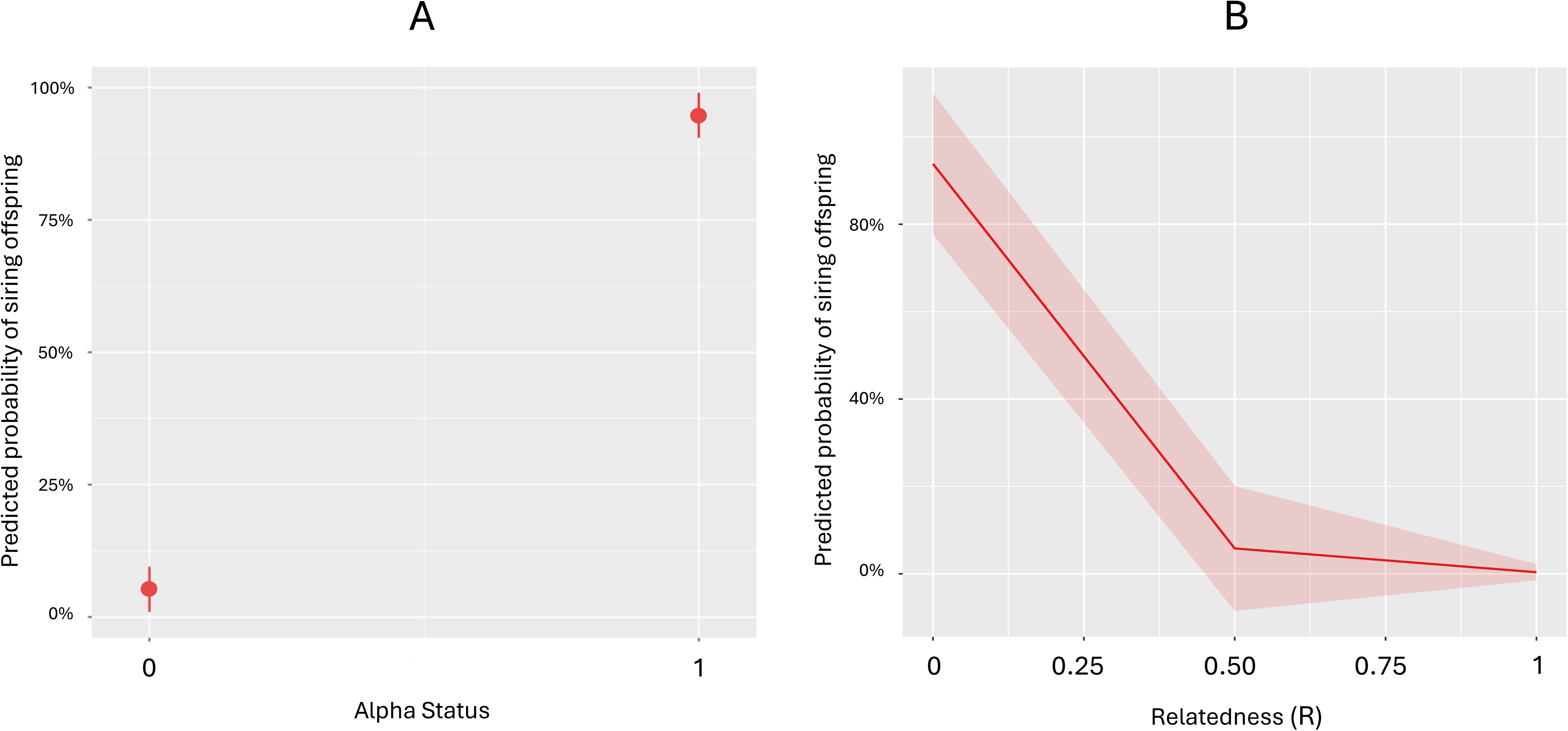
Predicted probabilities of a male siring an infant based on his (A) alpha status and (B) relatedness to the mother. Alpha males have a much higher probability of siring offspring than subordinate males. Unrelated males have a much higher probability of siring a female’s offspring.

**Table 2.**
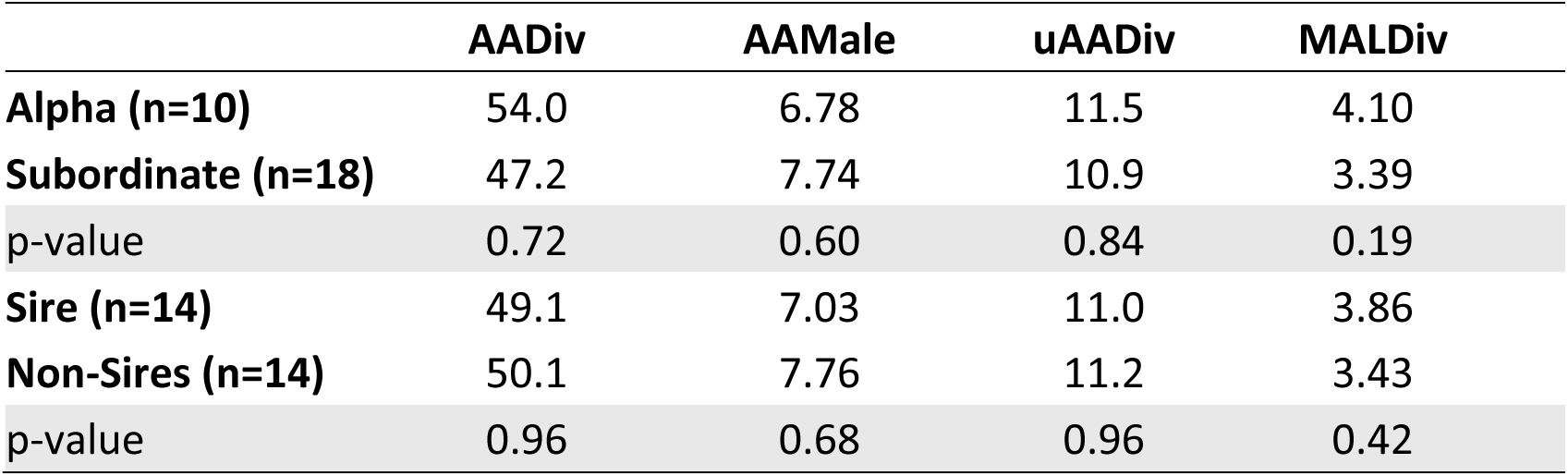
Results from t-tests comparing average MHC-DRB exon 2 values for male (N=28) metrics between alphas and subordinates, then sires and non-sires. Definitions of variables are included in Table 1. Note that individual males are counted as alpha if they ever attained alpha status, and as sire if they ever sired offspring in the study.

### MHC metrics do not predict the probability of siring offspring, or becoming alpha

After the correlation tests, we retained only one of the correlated metrics (MALDiv, MALDis, CALDis, and AAMale) for our analyses of MHC-DRB’s power to predict the probability of siring offspring. As expected, the *mclogit* models reveal that social status predicts siring success, where alpha males are more likely to be sires than subordinate males (Table 3; Fig. 1a). We also recovered relatedness as a statistically significant predictor of the likelihood of siring an infant (Table 3; Fig. 1b), in that the more closely related a potential mating pair is, the less likely the male is to sire offspring with that female. MHC variability at DRB exon 2 does not reliably predict the likelihood of males siring offspring in this population sample (Table 3). Additionally, MHC metrics do not reliably predict a male’s probability of becoming alpha (Table 4). These results are consistent across all four exons and the combined dataset (Table S2, S3).

**Table 3.**
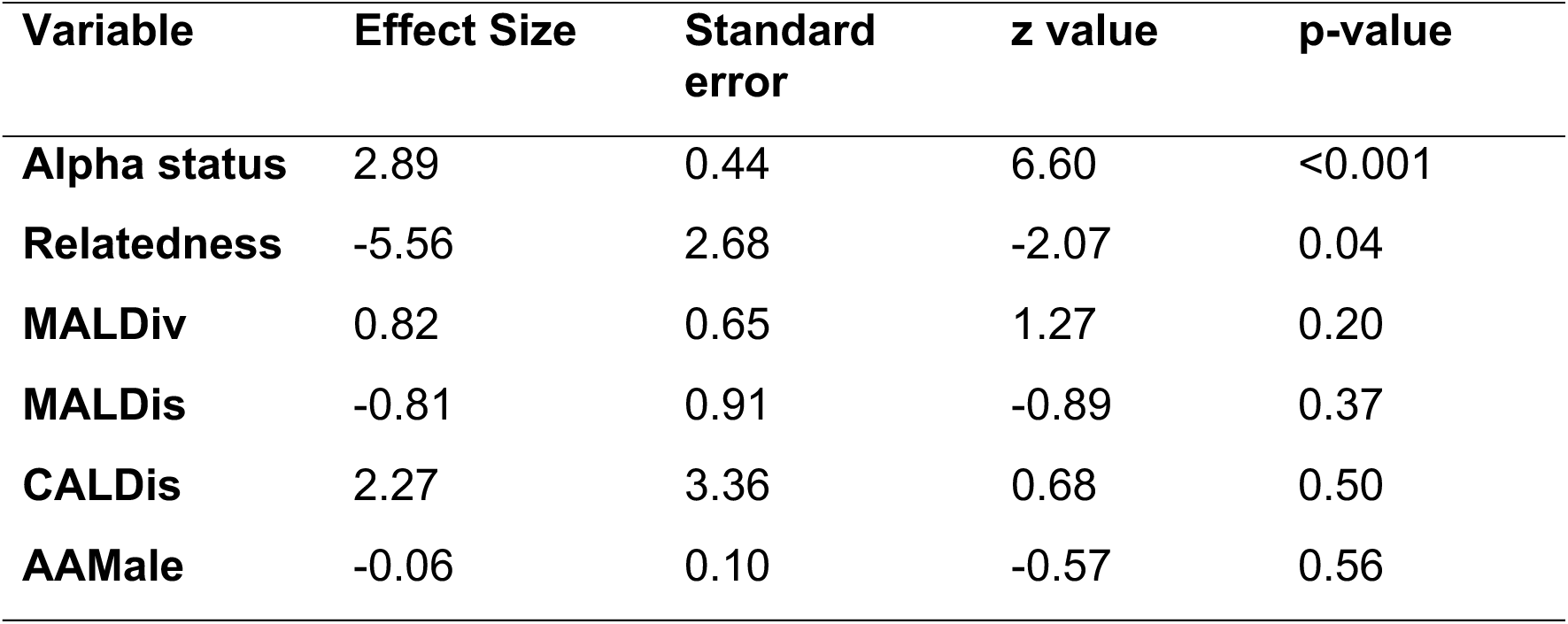
Result form fitting multinomial logit models with random effects for MHC DRB exon 2 to test if MHC metrics reliably predict siring success. See definitions in Table 1. Results were consistent when MHC varaiables were modeled independently to increase the number of observations/offspring (N) included in the analysis (not shown). We included only one metric when there was ≥ 0.8 positive pairwise correlations between metrics.

| Variable | Effect Size | Standard error | z value | p-value |
| --- | --- | --- | --- | --- |
| **Alpha status** | 2.89 | 0.44 | 6.60 | <0.001 |
| **Relatedness** | -5.56 | 2.68 | -2.07 | 0.04 |
| **MALDiv** | 0.82 | 0.65 | 1.27 | 0.20 |
| **MALDis** | -0.81 | 0.91 | -0.89 | 0.37 |
| **CALDis** | 2.27 | 3.36 | 0.68 | 0.50 |
| **AAMale** | -0.06 | 0.10 | -0.57 | 0.56 |

**Table 4.** Results from fitting generalized linear models to MHC-DRB exon 2 data to test if MHC metrics reliably predict a male (N=28) ascending to alpha status. See definitions in Table 1.

|  | Estimate | Std. Error | z value | Pr(>\|z\|) |
| --- | --- | --- | --- | --- |
| **Intercept** | -24.08 | 4630 | -0.005 | 0.996 |
| **MALDiv** | 10.78 | 1814 | 0.006 | 0.995 |
| **AADiv** | 0.20 | 245 | 0.001 | 0.999 |
| **uAADiv** | -4.37 | 2152 | -0.002 | 0.998 |
| **AAMale** | 2.97 | 1370 | 0.002 | 0.998 |

### Subordinate males heterozygous at MHC loci sire significantly more offspring than homozygous subordinates

While analyzing the data, we noted that relative to the number of offspring sired, subordinate males heterozygous at MHC loci sired nearly three times more offspring than MHC homozygotes (Figure 2). A z-test revealed a significant difference in proportions of successful sires between homozygous and heterozygous subordinate males (p=0.02). We therefore fit an *mclogit* model to only offspring sired by subordinate males for the MHC dataset. Consistent with previous results, MHC metrics did not predict likelihood of subordinates siring offspring at any of the individual loci or the combined dataset, although MALDiv and AAMale were nearly significant for DRB exon 3 (p ≈ 0.1; Table S4).We also found no significant differences in mean MHC metrics between subordinates who sired and those that did not (Table S5).

**Figure 2.**
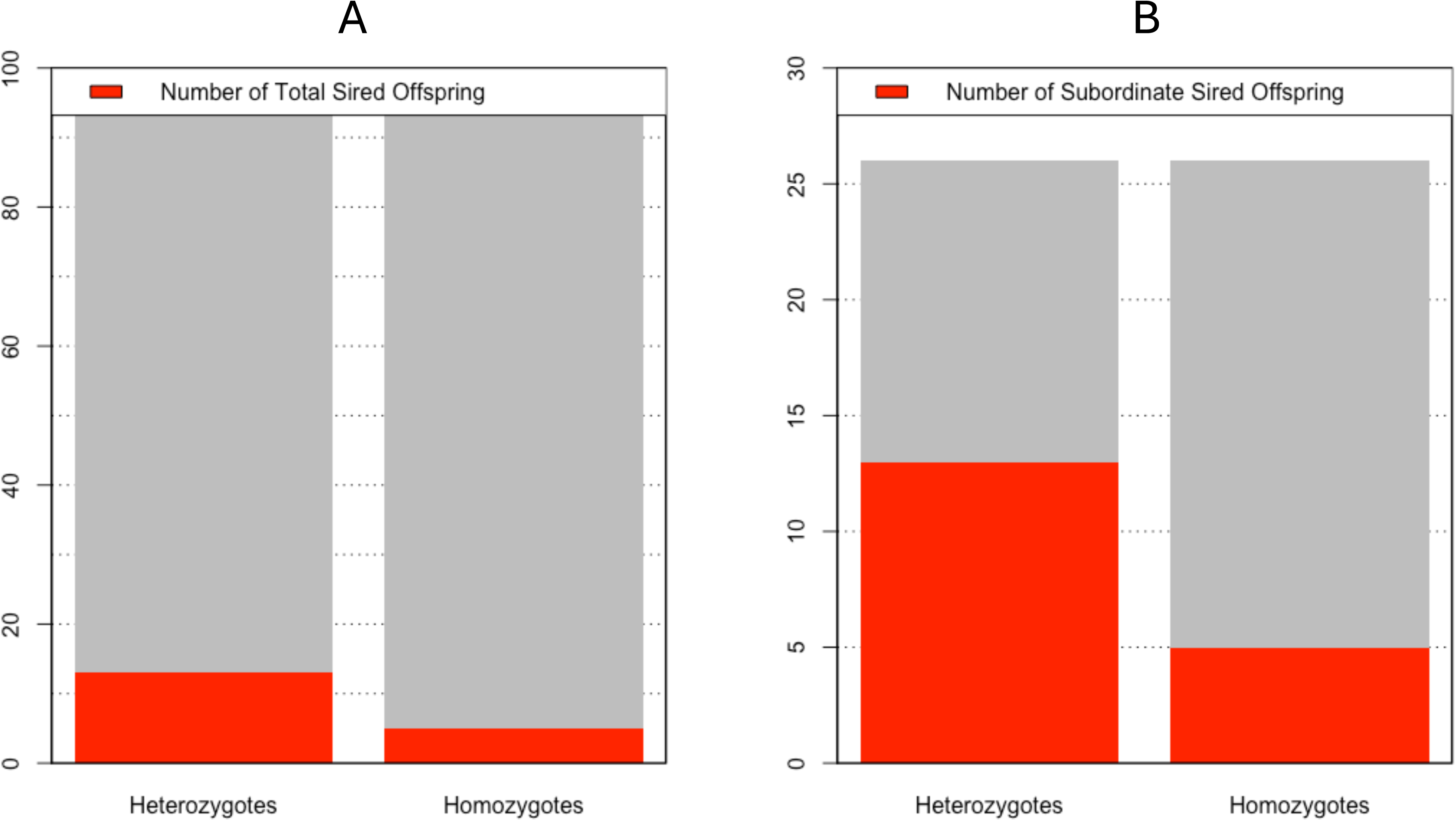
(A) Proportion of total offspring in the study (N=95) that were sired by MHC heterozygote or homozygote subordinate males. (B) Proportion of offspring sired by MHC heterozygous versus homozygous subordinate males (N=26). In both cases, these proportions are significantly different according to a z-test of proportion (p=0.02 and p=0.01, respectively).

### Offspring are more frequently heterozygous at MHC loci than expected

Based on the expected proportion of MHC-heterozygous offspring produced given the genotypes of the mothers and fathers, the proportion of expected heterozygous offspring was 0.59, and the observed heterozygosity was 0.68 (26 out of 38 offspring). In one-sided binomial simulations this was in the 90^th^ percentile (p = 0.155). However, when considering the probability of a female having a MHC-heterozygous offspring given the MHC genotypes of all potential sires in the group during each conception event, and taking the average probability of MHC heterozygous offspring across all recorded conception events with two or more genotyped sires available in the group, the proportion of expected MHC heterozygous offspring was 0.499, and the observed proportion of heterozygous offspring was 0.71 (27 of 38). In the one-sided binomial simulations this was in the 97.5% quartile and a significantly higher proportion of heterozygous offspring than expected by chance (p = 0.001).

## Discussion

We investigated the role of MHC genotypes in female mate choice and male reproductive success for SSR white-faced capuchin monkeys. With the high reproductive skew typical of this species, it might be expected that there is little opportunity for MHC genotypes to play a role in mate selection, unless genotypes impact the likelihood of becoming an alpha male. We found no statistically significant evidence that MHC genotypes influence the probability of an individual obtaining alpha status. We also found limited evidence for the MHC-diverse mate choice hypothesis and no evidence for the disassortative mate choice hypothesis.

We found some support for the MHC-diverse mate choice hypothesis among subordinate males that father more offspring in instances of inbreeding avoidance between the alpha male and his breeding daughters (Fig. 2). Thus, females may choose mates based on MHC heterozygosity when they produce offspring with subordinate males. Relatedly, we found that female mate choice led to a significantly higher proportion of MHC heterozygous offspring than predicted by chance given the mating pool. Although group living and reproductive skew in white-faced capuchins contrasts the solitary behavior in grey mouse lemurs, both species demonstrate an interplay between MHC-based mating decisions and inbreeding avoidance (Huchard et. al., 2013). Relative to the former, the disassortative mating at MHC-DRB in grey mouse lemurs may lead to higher probabilities of MHC-diverse offspring, while reduced genetic diversity in white-faced capuchins may mean that heterozygous subordinate males offer the best opportunity for MHC-diverse offspring when the female is not producing offspring with the alpha male. A formal study of overdominant selection (or heterozygote advantage) related to fitness in free-ranging rhesus macaques also found evidence for MHC-DQB heterozygous males experiencing more reproductive success (Sauerman et al., 2001). In the only other MHC-based mate choice study of a platyrrhine species, female muriquis also produced infants with males who were more MHC diverse than expected by chance, but not more MHC dissimilar to them (Chaves et al., 2023).

We may only find this suggestive evidence for MHC-mediated mate choice in white-faced capuchin monkeys because of their low population genomic diversity (Orkin et al., 2021). Although some studies show that patterns of MHC diversity may be decoupled from genome-wide variation in primates (Petersen et. al., 2022; Chaves et al., 2023), low MHC diversity in white-faced capuchins may reflect the strong effects of broader population dynamics. There are a few rare alleles, but across the four sampled groups in this population, there is a limited set of dominant haplotypes that combine to produce the observed genotypes in the parents and offspring sampled (Table 5; Buckner et al., 2021). Figure S1 shows the MHC DRB2 genealogy for the two largest main study groups in SSR; clearly the A and B haplotypes dominate the population. Our previous study of 70 adult individuals from four different social groups recovered only ten unique MHC DRB2 alleles, and individuals possessed two to six alleles (Buckner et al., 2021). By contrast, amongst 59 adult muriquis sampled and genotyped from a single social group, there were 22 unique MHC DRB2 alleles recovered, and individuals possessed three to eight MHC alleles (Chaves et al., 2023). Comparing DRB2 exon diversity across the two data sets, we see that muriquis have more genetic polymorphisms and more loci with amino acid changes than do the capuchins (Fig. 3), and the authors found significant support for MHC-diverse mate choice. Although additional factors may explain the contrasting results (e.g., hierarchical vs. egalitarian societies), a lack of variation in the sample set may limit the statistical power of mixed conditional logit models to detect signals in capuchins.

**Figure 3.**
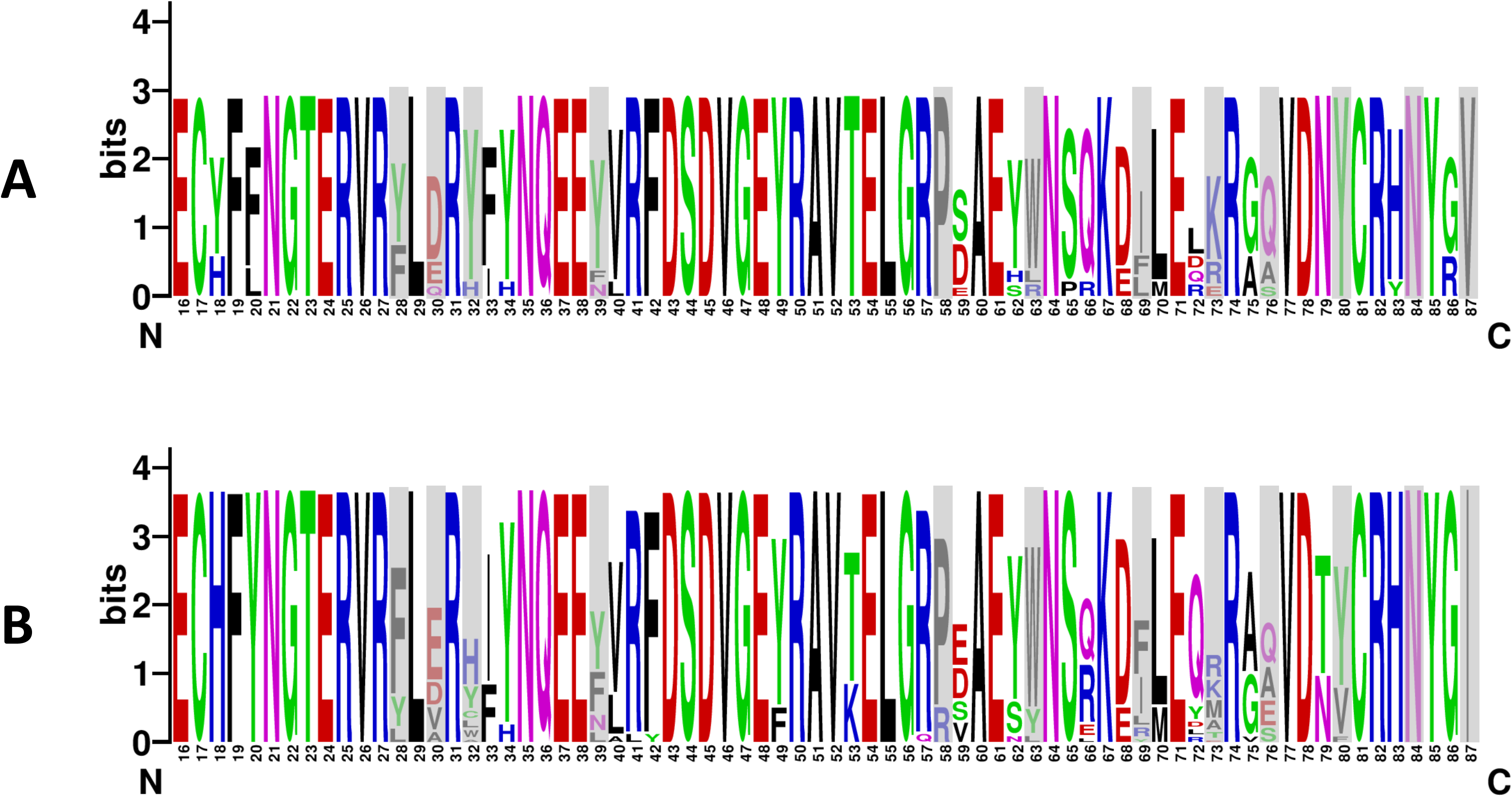
Sequence logos summarizing the amino acid diversity of DRB exon 2 for a population of (A) *Cebus imitator* (Buckner et al., 2021) and (B) *Brachyteles hypoxanthus* (Chaves et al., 2023). Amino acids are indicated by their single letter codes and the code colors indicate their major biochemical properties: Red–acidic; Blue–basic; Green– polar; Black–hydrophobic. Gray bars highlight predicted peptide binding region (PBR) sites based on putative human PBR (Reche and Reinherz, 2003). For the 72 amino acid sites summarized here, 21 sites were polymorphic for *C. imitator* and 24 for *B. hypoxanthus* and the latter generally displayed more diversity at putative peptide binding region sites (gray bars). Generated using WebLogo (Crooks et al., 2004).

**Table 5.**
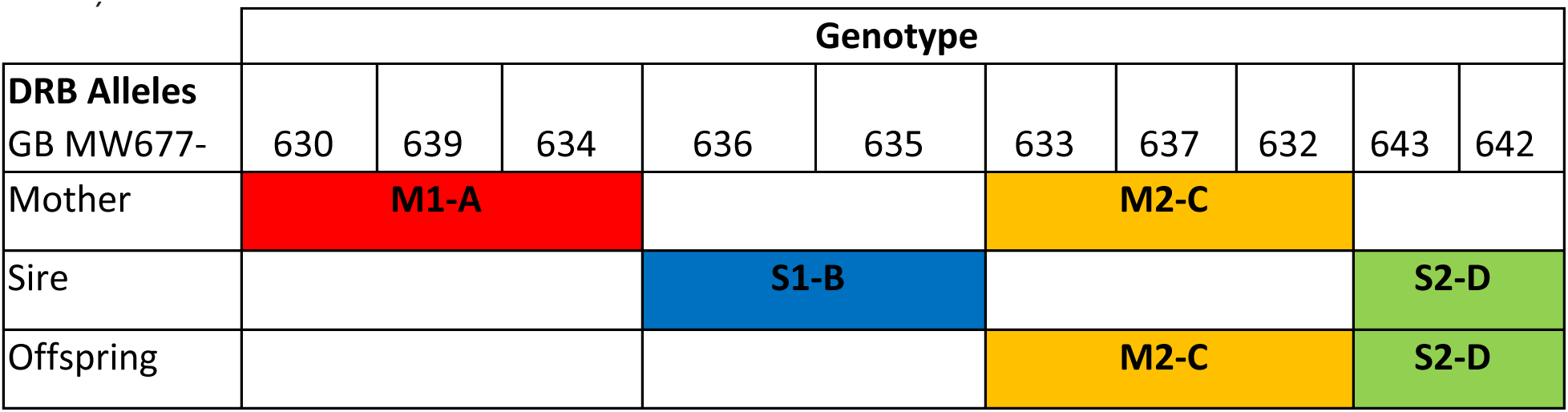
An example of an SSR *Cebus imitator* triad (mother, sire, and offspring) illustrating the four dominant haplotypes in the population (A, B, C, and D). M indicates the mother’s haplotypes, S indicates the sire’s haplotypes. The DRB exon 2 allele variants in each haplotype are listed using their genbank (GB) accession numbers (Buckner et al., 2021).

Additionally, the large effect sizes and large standard errors suggest issues of sample size and/or lack of variability in the sample, the latter perhaps reflecting the limited genotype diversity in the population. The high reproductive skew further reduces sample size as multiple observations (i.e., offspring births) involve various combinations of non-monogamous pairings of the same fathers and mothers and thus precludes the use of randomization tests to validate results (Hoover and Nevit, 2016). These qualities of the data also precluded tests of mate choice based on specific MHC alleles which may be more important than diversity or dissimilarity, and have been found in studies of other primates (e.g., chacma baboons - Huchard et al., 2010; mandrills - Setchell et al., 2009; fat-tailed dwarf lemurs [*Chierogaleus medius*]– Schwensow et al., 2008a; golden snub-nosed monkeys – Zhang et al., 2020). The difficulty of generating large sample sizes for wild primate populations remains a challenge for understanding the relationship between genetics and sexual selection. We hope that as long-term studies continue at sites like Santa Rosa, more comprehensive, systematic, non-invasive sampling of individuals across populations will provide more data to better inform analyses that test hypotheses about the role of genetics in mate choice, especially for species with complex social structures.

This study demonstrates the importance of accounting for the multiple factors that influence mate choice when testing for specific influences. As in humans, aspects of socio-cultural behavior may override, mitigate, or inhibit the ability to rely on signals of MHC genotype in mate choice (Dandine-Roulland, et. al., 2019) or may obfuscate the genetic patterns that could influence, or be influenced by mate choice. The range of results from MHC-associated mate choice studies in primates with similar social structure and mating strategies suggests that these social characteristics alone may not reliably predict the presence of MHC-based non-random mating. Ultimately, it may be most valuable to consider how patterns of selection for mate choice based on MHC diversity or dissimilarity may appear different based on the relative factors involved in disparate lineages (e.g., population dynamics, parasite loads, longevity, sociality, etc.). In different species, the relative and combined influence of certain behavioral, genetic, and ecological factors will likely result in distinct selective outcomes given the idiosyncratic nature of biological processes. We suspect that the factors themselves may be consistent across species, but the selective magnitude of the factors can change through phylogeny, space, and time. Understanding how social factors versus environmental factors and population history affect MHC variation will only be possible through more comparative studies. We suggest it would be most useful to analyze MHC-associated mate choice in several primate species at the same location (to control to some extent for ecology and land use change or isolation), as well as multiple populations of the same species, to help tease apart the multiple factors influencing the effects of MHC on mate choice.

## Conclusions

The role of MHC in mate choice in nonhuman primates remains under explored. To our knowledge, eight studies of MHC-based mate choice in wild primates have been undertaken (Schwensow et al., 2008a; Schwensow et al., 2008b; Huchard et al., 2010; Huchard et al., 2013; Yang et al., 2013; Zhang et al., 2020; Zhang et al., 2023; Chaves et. al., 2023). This study is only the second investigating MHC-mediated mate choice in a wild platyrrhine. Platyrrhines represent the most recent major geographical expansion of living primates, which likely resulted in a novel set of biological challenges, including exposure to new pathogens that would influence selection on the MHC. The extent to which sexual selection may have played a role in maintaining effective adaptive immunity during this transition is unclear without an understanding of the role of mate choice in platyrrhine MHC diversity and molecular evolution.

Alpha male status and male relatedness to the female best predicted siring success in this study, suggesting that female mate choice is either constrained by reproductive monopolization of the alpha male or that females prefer to mate with unrelated alpha males over subordinate males. While we found no evidence that MHC metrics are correlated with the probability of siring success in white-faced capuchins, we did find support for heterozygote advantage based on subordinate male siring success and offspring genotypes. Future studies will benefit from larger sample sizes that must necessarily come from long-term field studies with systematic sampling of individuals. We also acknowledge that the MHC locus is complex, and that different genes may show different patterns relative to mate choice so that more comprehensive characterization of the MHC (rather than reduced representation) may better illuminate the role of mate choice in maintaining MHC polymorphism and vice versa. With the advent of increasingly accurate sequencing technologies, reductions in per sample costs, and improvements to molecular methods addressing degraded DNA samples, we hope it will soon be practical to maximize the genetic information available from such samples to answer a variety of evolutionary questions.

## Acknowledgments

The authors thank the Costa Rican National Park Service and administrative team in Sector Santa Rosa of the Area de Conservacion Guanacaste (especially Roger Blanco Segura and Maria Marta Chavarria) for supporting our research and their assistance with permits and logistics. The authors would also like to thank the many people who contributed to the genetic sampling and life history data collection, especially R. Lopez, M. Myers, E. Walco, T. Busch, J. Rinderknecht, A.S. Pellier, A. Tecza, S. Millus, L. Wilkins, R. Jackson and K. Catanese. We would like to thank Andy Lin and Kotrina Kajokaite from UCLA Statistical Methods and Data Analytics for assistance with designing the analyses. The authors would like to acknowledge the following sources of funding for this research: Louisiana Board of Regents Support Fund, Research Competitiveness Subprogram (#077A-14) and Leakey Foundation to KMJ and JWL; Newcomb Institute, Stone Center for Latin American Studies, and the Lurcy and Lavin-Bernick Faculty Research Funds at Tulane University to KMJ; Leakey Foundation, National Science Foundation (#0926039), National Geographic Society (#8652-09), Natural Sciences and Engineering Research Council of Canada to VMS; and Natural Sciences and Research Council of Canada and Canada Research Chairs program to ADM and LMF.

