## Supplemental Tables and Figures for "MHC heterozygosity may increase subordinate but not alpha male siring success in white-faced capuchin monkeys (*Cebus imitator*)"

Table S1. Results from t-tests comparing mean MHC values for male (N=28) variables between alphas and subordinates, then sires and non-sires. Definitions of variables are included in Table 1.

| | AADiv | AAMale | uAADiv | MALDiv |
|---|---|---|---|---|
| | | **DRB exon 2** | | |
| **Alpha (n=10)** | 54.0 | 6.78 | 11.5 | 4.10 |
| **Subordinate (n=18)** | 47.2 | 7.74 | 10.9 | 3.39 |
| p-value | 0.72 | 0.60 | 0.84 | 0.19 |
| **Sire (n=14)** | 49.1 | 7.03 | 11.0 | 3.86 |
| **Non-Sires (n=14)** | 50.1 | 7.76 | 11.2 | 3.43 |
| p-value | 0.96 | 0.68 | 0.96 | 0.42 |
| | | **DRB exon 3** | | |
| **Alpha (n=10)** | 176 | 6.79 | 21.9 | 7.70 |
| **Subordinate (n=18)** | 130 | 7.50 | 19.2 | 6.44 |
| p-value | 0.09 | 0.08 | 0.09 | 0.11 |
| **Sire (n=14)** | 159 | 7.07 | 21.0 | 7.21 |
| **Non-Sires (n=14)** | 134 | 7.43 | 19.4 | 6.57 |
| p-value | 0.34 | 0.37 | 0.31 | 0.40 |
| | | **DQA** | | |
| **Alpha (n=10)** | 15.1 | 3.05 | 4.04 | 3.70 |
| **Subordinate (n=18)** | 14.1 | 3.63 | 4.17 | 3.33 |
| p-value | 0.60 | 0.05 | 0.73 | 0.06 |
| **Sire (n=14)** | 14.4 | 3.19 | 4.00 | 3.57 |
| **Non-Sires (n=14)** | 14.5 | 3.65 | 4.24 | 3.36 |
| p-value | 0.97 | 0.11 | 0.48 | 0.27 |
| | | **DQB** | | |
| **Alpha (n=10)** | 39.7 | 7.98 | 10.6 | 3.70 |
| **Subordinate (n=18)** | 35.2 | 8.41 | 10.0 | 3.44 |
| p-value | 0.28 | 0.24 | 0.22 | 0.27 |
| **Sire (n=14)** | 38.6 | 8.06 | 10.5 | 3.64 |
| **Non-Sires (n=14)** | 35.0 | 8.46 | 9.98 | 3.43 |
| p-value | 0.36 | 0.25 | 0.30 | 0.33 |
| | | **Combined** | | |
| **Alpha (n=10)** | 284 | 24.6 | 48.1 | 19.2 |
| **Subordinate (n=18)** | 227 | 27.3 | 44.3 | 16.6 |
| p-value | 0.22 | 0.17 | 0.45 | 0.13 |

| | | | | |
|---|---|---|---|---|
| **Sire (n=14)** | 261 | 25.3 | 46.5 | 18.3 |
| **Non-Sires (n=14)** | 233 | 27.3 | 44.8 | 16.8 |
| p-value | 0.54 | 0.30 | 0.73 | 0.37 |

Table S2. Results from fitting generalized linear models to MHC data variables to test if these variables reliably predict a male (N=28) ascending to alpha status. Definitions of variables are included in Table 1. Values for AAMale in DQB exon 3 were not defined because this metric had an exact linear relationship with another included variable.

| | Estimate | Std. Error | z value | Pr(>\|z\|) |
|---|---|---|---|---|
| | | **DRB exon 2** | | |
| **Intercept** | -24.1 | 4630 | -0.005 | 1.00 |
| **MALDiv** | 10.8 | 1814 | 0.006 | 0.99 |
| **AADiv** | 0.20 | 245 | 0.001 | 1.00 |
| **uAADiv** | -4.37 | 2152 | -0.002 | 1.00 |
| **AAMale** | 2.97 | 1370 | 0.002 | 1.00 |
| | | **DRB exon 3** | | |
| **Intercept** | 83.8 | 67.3 | 1.24 | 0.21 |
| **MALDiv** | -7.76 | 6.79 | -1.14 | 0.25 |
| **AADiv** | 0.26 | 0.28 | 0.92 | 0.36 |
| **uAADiv** | -1.73 | 2.46 | -0.70 | 0.48 |
| **AAMale** | -4.71 | 3.93 | -1.20 | 0.23 |
| | | **DQA exon 3** | | |
| **Intercept** | -9.48 | 2.73e5 | 0 | 1.00 |
| **MALDiv** | 4.25 | 9.11e4 | 0 | 1.00 |
| **AADiv** | -4.15e8 | 2.24e9 | -0.18 | 0.85 |
| **UAADiv** | 2.49e9 | 1.35e10 | 0.18 | 0.85 |
| **AAMale** | 1.25e9 | 6.73e9 | -0.18 | 0.85 |
| | | **DQB exon 3** | | |
| **Intercept** | -72.40 | 6588 | -0.01 | 0.99 |
| **MALDiv** | 22.93 | 2196 | 0.01 | 0.99 |
| **AADiv** | -2.01 | 195 | -0.01 | 0.99 |
| **UAADiv** | 6.34 | 585 | 0.01 | 0.99 |

| | | | | |
|---|---|---|---|---|
| | | **Combined** | | |
| **Intercept** | -854 | 70131 | -0.01 | 0.99 |
| **MALDiv** | 96.6 | 7932 | 0.01 | 0.99 |
| **AADiv** | 3.76 | 314 | 0.01 | 0.99 |
| **UAADiv** | -71.8 | 5939 | -0.01 | 0.99 |
| **AAMale** | 56.7 | 4694 | 0.01 | 0.99 |

Table S3. Results from fitting multinomial logit models with random effects for MHC data to test if MHC metrics reliably predict siring success. Results were consistent when MHC variables were modeled independently to increase the number of observations/offspring (N) included in the analysis (not shown). Correlations between variables were calculated separately for each dataset, so variables evaluated for each locus differ slightly. Definitions of variables are included in Table 1.

| Variable | Effect Size | Standard error | z value | p-value |
|---|---|---|---|---|
| | | **DRB exon 2 (N = 74)** | | |
| Alpha status | 2.89 | 0.44 | 6.60 | 4.06e-11 |
| Relatedness | -5.56 | 2.68 | -2.07 | 0.04 |
| MALDiv | 0.82 | 0.65 | 1.27 | 0.20 |
| MALDis | -0.81 | 0.91 | -0.89 | 0.37 |
| CALDis | 2.27 | 3.36 | 0.68 | 0.50 |
| AAMale | -0.06 | 0.10 | -0.57 | 0.56 |
| | | **DRB exon 3 (N = 73)** | | |
| Alpha status | 3.08 | 0.46 | 6.68 | 2.36e-11 |
| Relatedness | -6.55 | 2.64 | -2.48 | 0.01 |
| MALDiv | 0.46 | 0.42 | 1.11 | 0.27 |
| MALDis | -0.60 | 0.66 | -0.90 | 0.37 |
| CALDis | 2.88 | 4.02 | 0.72 | 0.47 |
| | | **DQA exon 3 (N = 73)** | | |
| Alpha status | 2.86 | 0.42 | 6.77 | 1.28e-11 |
| Relatedness | -5.31 | 2.60 | -2.05 | 0.04 |
| MALDiv | 3.60 | 3.09 | 1.16 | 0.24 |
| MALDis | -5.07 | 5.27 | -0.96 | 0.33 |
| CALDis | 11.21 | 12.42 | 0.90 | 0.37 |
| | | **DQB exon 3 (N = 74)** | | |
| Alpha status | 2.94 | 0.42 | 6.93 | 4.21e-12 |
| Relatedness | -6.22 | 2.31 | -2.70 | 0.007 |
| MALDiv | 1.96 | 1.62 | 1.21 | 0.22 |

| | | | | |
|---|---|---|---|---|
| MALDis | -1.73 | 2.48 | -0.70 | 0.48 |
| CALDis | 2.46 | 6.06 | 0.41 | 0.68 |
| **Combined (N = 72)** | | | | |
| Alpha status | 3.04 | 0.46 | 6.64 | 3.14e-11 |
| Relatedness | -6.46 | 2.59 | -2.50 | 0.01 |
| MALDiv | 0.25 | 0.22 | 1.12 | 0.26 |
| MALDis | -0.30 | 0.35 | -0.87 | 0.38 |
| CALDis | 0.95 | 1.37 | 0.70 | 0.49 |

Table S4. Results from fitting multinomial logit models with random effects to a subset of MHC data including only offspring sired by subordinate males. Sample sizes may be too small to estimate reliable effect sizes. Definitions of variables are included in Table 1.

| Variable | Effect size | Standard Error | z value | p-value |
|---|---|---|---|---|
| | | **DRB exon 2 (N=14)** | | |
| Relatedness | 2.01 | 1.55 | 1.30 | 0.19 |
| MALDiv | -2.87 | 2.61 | -1.10 | 0.27 |
| MALDis | 8.65 | 6.99 | 1.24 | 0.21 |
| CALDis | -0.05 | 0.13 | -0.38 | 0.70 |
| AAMale | 2.01 | 1.55 | 1.30 | 0.19 |
| | | **DRB exon 3 (N = 13)** | | |
| Relatedness | -9.60 | 6.98 | -1.38 | 0.17 |
| MALDiv | 19.2 | 11.8 | 1.63 | 0.10 |
| MALDis | -2.40 | 2.13 | -1.13 | 0.26 |
| CALDis | 13.00 | 10.17 | 1.27 | 0.20 |
| AAMale | 35.7 | 22.54 | 1.59 | 0.11 |
| | | **DQA exon 3 (N = 14)** | | |
| Relatedness | -2.9 | 5.29 | -0.55 | 0.579 |
| MALDiv | 4.33 | 6.06 | 0.71 | 0.475 |
| MALDis | -8.45 | 11.0 | -0.76 | 0.444 |
| CALDis | 20.0 | 23.0 | 0.87 | 0.384 |
| AAMale | -0.79 | 0.93 | -0.85 | 0.395 |
| | | **DQB exon 3 (N = 14)** | | |
| Relatedness | -5.1771 | 4.4652 | -1.159 | 0.246 |
| MALDiv | -2.9120 | 3.6671 | -0.794 | 0.427 |
| MALDis | 0.4468 | 4.1021 | 0.109 | 0.913 |
| CALDis | -0.6390 | 8.9354 | -0.072 | 0.943 |
| AAMale | -2.4969 | 2.3140 | -1.079 | 0.281 |
| | | **Combined (N = 13)** | | |

| | | | | |
|---|---|---|---|---|
| Relatedness | -3.33271 | 5.45898 | -0.611 | 0.542 |
| MALDiv | 0.68810 | 0.71535 | 0.962 | 0.336 |
| MALDis | -1.18778 | 1.29199 | -0.919 | 0.358 |
| CALDis | 3.84972 | 4.01034 | 0.960 | 0.337 |
| AAMale | -0.01112 | 0.12605 | -0.088 | 0.930 |

Table S5. Results from t-test for MHC data for subordinate sires vs non-sires (exon 3 of DQA, DQB and DRB, exon 2 of DRB, and summed combined data). Definitions of other variables are included in Table 1. This test includes all males that sired offspring as subordinates, even if they later ascended to alpha. Results were consistent when males that were both alphas and subordinates were removed from the analysis (not shown).

| | AADiv | AAMale | uAADiv | MALDiv |
|---|---|---|---|---|
| | **DRB exon 2** | | | |
| **Sires (n=8)** | 48.0 | 7.01 | 10.9 | 3.75 |
| **Non-Sires (n=17)** | 50.5 | 7.42 | 11.1 | 3.53 |
| p-value | 0.91 | 0.84 | 0.95 | 0.72 |
| | **DRB exon 3** | | | |
| **Sires (n=8)** | 153.75 | 7.25 | 20.75 | 7.00 |
| **Non-Sires (n=17)** | 141.65 | 7.33 | 19.90 | 6.76 |
| p-value | 0.69 | 0.85 | 0.64 | 0.79 |
| | **DQA exon 3** | | | |
| **Sires (n=8)** | 14.0 | 3.25 | 3.96 | 3.50 |
| **Non-Sires (n=17)** | 14.5 | 3.54 | 4.19 | 3.41 |
| p-value | 0.80 | 0.40 | 0.58 | 0.69 |
| | **DQB exon 3** | | | |
| **Sires (n=8)** | 38.37 | 8.10 | 10.4 | 3.62 |
| **Non-Sires (n=17)** | 35.22 | 8.41 | 10.0 | 3.44 |
| p-value | 0.50 | 0.44 | 0.42 | 0.48 |
| | **Combined** | | | |
| **Sires (n=8)** | 254 | 25.6 | 46.0 | 17.87 |
| **Non-Sires (n=17)** | 242 | 26.7 | 45.3 | 17.18 |
| p-value | 0.83 | 0.62 | 0.90 | 0.72 |

Supplemental Figure 1. MHC DRB exon 2 matrilineal genealogy for the two largest study groups in SSR. For each group, the first row indicates matriarchs of groups, and for subsequent rows, the first line of each row indicates genotypes of each offspring of that female, the second line indicates the genotype of the sire with whom the female produced those offspring. A, B, C and D haplotypes correspond to A, B, C and D allele sets described in Table 5.

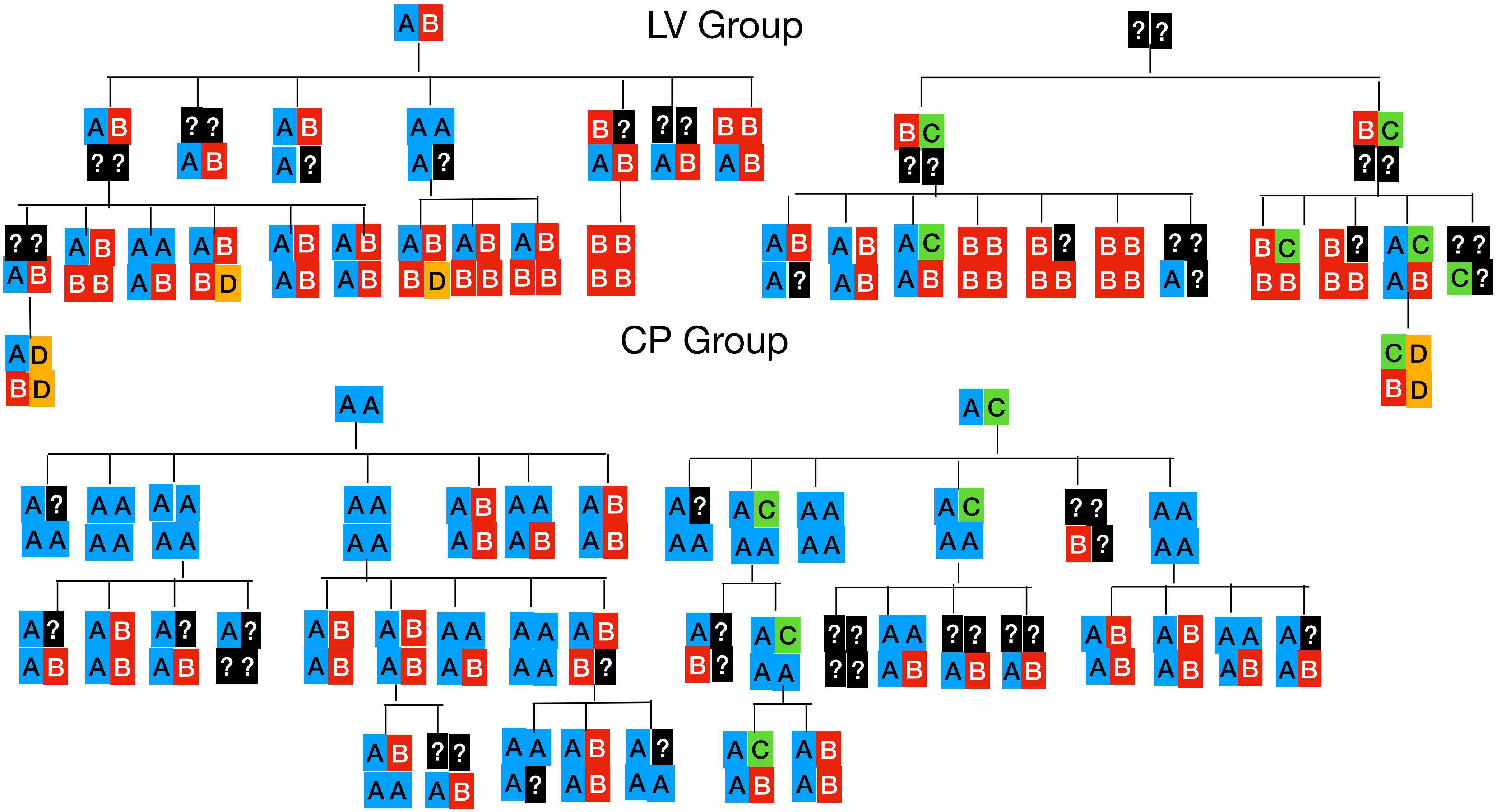
LV Group
CP Group
